# Individual stochasticity in the life history strategies of animals and plants

**DOI:** 10.1101/2022.03.06.483187

**Authors:** Pablo José Varas-Enríquez, Silke van Daalen, Hal Caswell

## Abstract

The life histories of organisms are expressed as rates of development, reproduction, and survival. However, individuals may experience differential outcomes for the same set of rates. Such individual stochasticity generates variance around familiar mean measures of life history traits, such as life expectancy and the reproductive number *R*_0_. By writing life cycles as Markov chains, we calculate variance and other indices of variability for longevity, lifetime reproductive output (LRO), age at offspring production, and age at maturity for 83 animal and 332 plant populations from the Comadre and Compadre matrix databases. We find that the magnitude within and variability between populations in variance indices in LRO, especially, are surprisingly high. We furthermore use principal components analysis to assess how the inclusion of variance indices affects life history constraints. We find that these indices, to a similar or greater degree than the mean, explain the variation in life history strategies among plants and animals.

## 1 Introduction

The life history of any organism can be described as movement among a set of stages, with individuals moving at rates that depend on the phenotype and the environment. The stages may be defined in terms of age, size, developmental stage, or other criteria. Any demographic model for the life history includes at least three processes: development (transitions among the living stages), mortality (transition out of the set of living stages), and fertility (production of new individuals). Each of these processes is stochastic. Mortality or survival happen with a probability. Reproduction, and the number and kind of offspring produced, are stochastic. Transitions during the developmental process are equally governed by probabilities.

Stochastic transitions lead to variation among individuals in longevity, reproduction, and development, even if they experience identical rates and risks at every stage. This variation is said to be due to individual stochasticity (Caswell, 2009). It is critically important to distinguish the variation resulting from individual stochasticity from variation due to heterogeneity among individuals. A demographic model describes the rates applying to all individuals in a given stage. Heterogeneity describes variation due to differences among individuals in the vital rates they experience in a given stage. Calculations of variance from a demographic model explicitly assume that all individuals experience the same rates at any life cycle stage, and variances resulting from individual stochasticity provide a critical standard of comparison in the search for evidence of heterogeneity (e.g., Steiner et al., 2010; Tuljapurkar et al., 2009; van Daalen & Caswell, 2017; Hartemink et al., 2017).

Population biology and human demography have long focused on expected values of the outcomes of demographic models, and these expected values are usually considered as life history traits. Life expectancy, for example, is the expectation of longevity; the net reproductive rate *R*_0_ is the expected value of lifetime reproduction; and the generation time is the mean age at reproduction. But expected values give only part of the story, and it is now possible to calculate the variation, due to individual stochasticity in the life cycle, of many life history outcomes.

The variability implied by a life cycle is calculated by embedding the population projection matrix into a finite state, discrete time, absorbing Markov chain. Such a chain includes transient states corresponding to living stages of the life cycle, and one or more absorbing states corresponding to death or other ways of leaving the living states. Longevity is the time required to reach an absorbing state; its mean, variance, and other moments are calculated from the Markov chain. Lifetime reproductive output is calculated by associating a ‘reward’ with every transition in the Markov chain. The reward represents stage-specific reproductive output; the Markov chain provides the mean variance, skewness, and all the moments of lifetime reproductive output, starting from any living state. For examples of Markov chain methods applied to individual stochasticity (or what would now be called individual stochasticity), see Cochran & Ellner (1992), Caswell (2001, 2009, 2011), Steinsaltz & Evans (2004), Tuljapurkar & Horvitz (2006); Tuljapurkar et al. (2009), Horvitz & Tuljapurkar (2008), Steiner & Tuljapurkar (2012), van Daalen & Caswell (2015, 2017).

Quantifying the variance in demographic outcomes is important because any analysis of a random variable is incomplete until it includes some indication of variation. In the case of life histories, variance due to heritable differences among individuals in fitness-related traits is potential raw material for natural selection. Variance due to individual stochasticity is not, so it is important to know how much variance is implied by the life cycle, as a null model for evaluating potential for selection (Steiner & Tuljapurkar, 2012). In applied contexts, planning and management require some quantification of risk (as actuaries and investors are well aware) and risk can be measured by variance, skewness and other measures of variability.

Because variation in demographic outcomes is implied by, and can be calculated from, a demographic model, it should be treated as a life history trait (van Daalen & Caswell, 2020a) and incorporated into comparative life history studies, just as the mean values are. This study provides the first detailed cross-taxon analysis of the variation in demographic outcomes, treated as life history traits, and how those measures of variation relate to other life history traits, across species.

We will consider individual stochasticity in a range of demographic outcomes (lifetime reproductive output, longevity, age at maturity, measures of iteroparity, and others). They will be calculated using Markov chains and Markov chains with rewards, and applied to population projection matrices obtained from the Comadre Animal Matrix Database and the Compadre Plant Matrix Database (Salguero-Gómez et al., 2015, 2016).

Our first goal is to quantify the amount of variation in demographic outcomes within populations for a large collection of plant and animal species. We will explore the distribution among populations of those measures of variation. Then we will examine how the measures of variation fit into the patterns of association among life histories, by extracting Principal Components that arrange species along life history axes. These axes can be interpreted in terms of combinations of life history traits, as has been done before for *r*- and *K*-selection (a binary contrast) (Pianka, 1970), the fast-slow continuum (a one-dimensional arrangement) (Stearns, 1983), and more recent two-dimensional arrangements (Salguero-Gómez et al., 2016). Our results reveal patterns that are not apparent in analyses restricted to mean properties.

The paper is organized as follows. In Section 2 we describe the sources of our demographic data, and in Section 3 we describe the calculation of the statistics of demographic outcomes. In Section 4 we report the patterns of variability for lifetime reproductive output and longevity, and in Section 5 we present the life history patterns emerging from a multidimensional approach; we end with a Discussion. Details of the calculations, and supplementary figures, are presented in Supporting Information.

## 2 Data

A comparative analysis of demographic outcomes requires demographic information for a diverse set of species. To obtain this information, we use the Comadre and Compadre matrix databases. These databases are open-data online repositories of high-quality, parameterized matrix population models for animal (Comadre v2.0.1) and plant (Compadre v4.0.1) species (Salguero-Gómez et al., 2015, 2016, obtained from www.compadre-db.org).

From these databases we selected those matrix population models that represented natural (i.e. unmanipulated) environments. We limited our analysis to matrices that were defined as an arithmetic average of matrices defined for a single time point (as provided by the database), with an annual projection interval, with non-zero fertility values, and with at least five stages. We eliminated two-sex models because they create life cycle pathways that do not appear in the more common single-sex models. We also eliminated matrices including clonal reproduction because they would require a separate analysis of lifetime reproductive output to separate clonal and sexual reproduction.

Calculations of lifetime demographic outcomes make frequent use of the fundamental matrix **N** = (**I** − **U**)^−1^. We excluded matrices for which the condition number of (**I** − **U**) *>* 1000; these matrices are near-singular (the species is near-immortal) and the fundamental matrix is unreliable.

Examining the matrices, we found that some life histories are modelled in a way that creates enormous values of the statistics of LRO (e.g., variances on the order of 10^11^). This often results from an unfortunate choice by the investigators of how to combine, within one life cycle, the production of huge numbers of seeds or larvae and a miniscule fraction of these probabilities that survive to enter the life cycle. To prevent these models from artifactually dominating all of our analytical results we applied a method of Kimber (1990) to screen for and remove outliers. The method is designed specifically for skewed data and identifies robust upper and lower bounds defined in terms of the quartiles of the data. Let *Q*_1_, *Q*_2_, and *Q*_3_ be the 0.25, 0.5, and 0.75 quantiles of the data. The upper limit defining outliers is *Q*_3_ + 2*c* (*Q*_3_ − *Q*_2_). After some experimentation on our part we chose *c* = 5 as a value that identified a group of genuine outliers. We excluded matrices for which the mean, variance, or kurtosis of LRO exceeded this value.

These criteria resulted in sets of 83 animal matrices and 332 plant matrices from the Comadre and Compadre databases respectively, representing 47 animal and 141 plant species.

## 3 Calculation of individual stochasticity in life history traits

Life history traits in common usage include those related to survival and longevity (e.g., life expectancy), to the amount of reproduction (e.g., *R*_0_, total fertility rate), and to the timing of reproduction and its distribution over the lifespan (e.g., age at maturity, generation time, iteroparity). Our approach differs because we will compute not only the mean of these traits, but also the higher moments and measures of variance, skewness, and kurtosis.

These results are obtained from the population matrices. The database supplies, for each population, a transition matrix **U** and a fertility matrix **F**. The (*i, j*) entry of **U** is the probability of transition from stage *j* to stage *i*, conditional on survival. The (*i, j*) entry of **F** is the mean number of stage *i* offspring produced by a stage *j* individual in a unit of time. The matrix **U** forms the basis of a finite-state, discrete-time, Markov chain. The life cycle contains *τ* transient (i.e., living) stages and *α* absorbing (i.e., dead or removed) stages. The transition probability matrix for the chain is

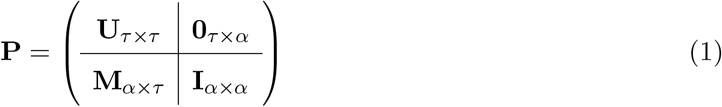

where the (*i, j*) entry of **M** is the transition probability from the *j*th living stage to the *i*th absorbing stage. The total number of states is *s* = *α* + *τ*.

From the Markov chain matrix **P** and the fertility matrix **F** we calculate life history traits as follows. Details of the calculation are given in the Supplementary Materials.

### Longevity

The longevity of an individual is the time required to leave the set of transient states (Caswell, 2001, 2009). A key piece in all the calculations is the fundamental matrix **N**, given by

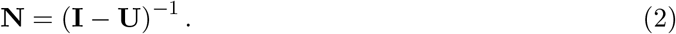

The (*i, j*) entry of **N** is the mean time spent in stage *i*, starting from stage *j*, before eventual absorption (death). We calculated the first four moments of longevity, starting from the first stage in the life cycle, using the expressions in equations (21)–(24) in Caswell (2013). From these moments we calculated the mean, standard deviation, coefficient of variation (CV), skewness, and kurtosis of longevity.

### Lifetime reproductive output (LRO)

We calculated the statistics of LRO using Markov chains with rewards (MCWR, Caswell, 2011; van Daalen & Caswell, 2015, 2017). This analysis treats reproduction as a reward collected by an individual at each step in its life, up until death. In the absence of data on the distribution of individual stage-specific reproduction, we used a Poisson distribution to create the reproductive reward matrices. For species that produce multiple types of offspring (i.e., in which **F** contains positive entries in more than one row), we summed all types at each age and treated the sum as the mean stage-specific reproductive output. We computed the first four moments of LRO according to van Daalen & Caswell (2017, Theorem 1). From these moments we calculated the mean, variance, standardized variance, skewness, and kurtosis of LRO. The standardized variance in LRO; i.e., the ratio of the variance to the square of the mean, is frequently calculated. Crow (1958) showed that the standardized variance in fitness is the rate at which selection would increase fitness if all the variance were genetic.

### Distribution of reproduction: generation time and iteroparity

Reproduction happens at different ages during the life of an individual. The distribution, over the lifetime, of reproduction provides two useful demographic outcomes: generation time and a measure of iteroparity.

The cohort generation time is the mean age at which offspring are produced by an individual. The formula for age-classified models is classical (e,g., Coale, 1972), but we need a version that applies also to stage-classified models. The necessary result is found in Caswell (2009, Appendix A); see also Caswell (2019, Chap. 5). When multiple types of offspring exist, we summed all types of offspring at every stage before calculating the generation time.

The distinction between semelparity and iteroparity (reproducing only once vs. being capable of reproducing repeatedly) focuses on the extent to which reproduction is spread over the life cycle. The degree of iteroparity is difficult to analyze demographically because of the rarity of models that explicitly include parity (see Caswell (2020) for an example of an age-parity model). We calculated an index of iteroparity by extending the calculation of generation time to compute the variance, and thence the coefficient of variation of the ages at which offspring are produced. Strict semelparity at a fixed age (as in the periodical cicadas *Magicicada sp*.) would have a CV of 0. If reproduction is spread over many ages, the CV will be greater than 0 and measure how widely spread the ages at reproduction are. The derivation of this measure of iteroparity is given in the Supplementary Material.

### Age at maturity

We calculated the mean, standard deviation, and coefficient of variation of the age at maturity using the methods of Caswell (2001, Section 5.3.3). This involves creating an absorbing set consisting of all reproductive stages, and calculating the moments of the time required to reach that absorbing set.

### Measures of uncertainty

The measurement of the variability of any trait requires some thought. In addition to the variance and related statistics, we calculated the skewness and kurtosis of longevity and of lifetime reproductive output, as measurements of the uncertainty of individual fates. Skewness is a measure of asymmetry of the distribution, and high values of skewness imply that the variable is even less certain than might be expected from the variance. Kurtosis measures the ‘heaviness’ of the tails of the distribution, and thus gives an indication of the likelihood of extreme values. The variance, skewness, and kurtosis were calculated from the moments using the formulas from Caswell (2011) and van Daalen & Caswell (2017).

## 4 Individual stochasticity generates variability: how much?

In this section we explore some of the patterns generated by individual stochasticity in the statistics of lifetime reproductive output and longevity. We focus on these patterns because they are important components of fitness. Longevity integrates the chances of survival over all the pathways in the life cycle. LRO integrates both survival and fertility. Our preliminary analysis showed that both of these quantities are highly variable, both within and between the populations in our subset.

Individual stochasticity creates a tremendous amount of variability in lifetime reproductive output in both animals and plants. Table 1 shows the median (preferable to the mean as a measure of central tendency for skewed data) and quantiles (2.5%, 25%, 75%, 97.5%) of each of the statistics of LRO and longevity. The 25% and 75% quantiles define the interquartile range, capturing the central 50% of the observations. The 2.5% and 97.5% quantiles capture 95% of the observations, and can be treated as the “range” of variation without the instability of minimum and maximum values. The scaled measures of variation (CV, standardized variance, skewness, kurtosis) are particularly useful indices for comparative purposes, because they are dimensionless.

**Table 1:** Quantiles of lifetime reproductive output (LRO) and longevity for the subsets of animals and plants.

| **LRO** | $Q_{2.5}$ | $Q_{25}$ | Median | $Q_{75}$ | $Q_{97.5}$ |
| --- | --- | --- | --- | --- | --- |
| Animals |  |  |  |  |  |
| Mean | 0.11 | 0.60 | 1.14 | 2.01 | 6.60 |
| Variance | 0.26 | 2.10 | 6.41 | 22.00 | 201.08 |
| OFS | 1.09 | 3.04 | 4.92 | 9.94 | 90.37 |
| Skewness | 1.00 | 2.35 | 3.37 | 4.45 | 13.19 |
| Kurtosis | 0.08 | 6.61 | 15.27 | 25.37 | 216.85 |
| Plants |  |  |  |  |  |
| Mean | 0.02 | 0.30 | 1.09 | 3.20 | 18.72 |
| Variance | 0.07 | 6.18 | 67.57 | 1416.0 | 38073 |
| OFS | 2.12 | 13.06 | 75.96 | 584.8 | 6486.9 |
| Skewness | 2.26 | 5.37 | 13.71 | 35.49 | 133.28 |
| Kurtosis | 6.92 | 38.86 | 244.4 | 1674.8 | 24087 |
| **Longevity** |  |  |  |  |  |
| Animals |  |  |  |  |  |
| Mean | 1.16 | 2.99 | 4.57 | 6.93 | 31.08 |
| St. Dev. | 0.71 | 3.35 | 5.38 | 8.57 | 28.03 |
| CV | 0.56 | 0.88 | 1.07 | 1.25 | 1.89 |
| Skewness | 0.34 | 1.99 | 2.31 | 3.08 | 9.01 |
| Kurtosis | -1.25 | 4.32 | 7.30 | 11.10 | 79.91 |
| Plants |  |  |  |  |  |
| Mean | 1.03 | 1.99 | 3.26 | 6.81 | 41.98 |
| St. Dev. | 0.50 | 2.25 | 5.15 | 11.76 | 105.53 |
| CV | 0.37 | 0.87 | 1.26 | 2.12 | 4.72 |
| Skewness | 0.66 | 2.34 | 4.31 | 9.16 | 53.71 |
| Kurtosis | 2.15 | 8.01 | 26.59 | 133.86 | 3852.7 |

### 4.1 Lifetime reproductive output

In animals, the median standardized variance (opportunity for selection) in LRO is 4.92, and is typically (i.e., 50% of the cases) between 3.04 and 9.94. In plants, the median is 75.96, with typical values ranging from 13.06 to 584.8. LRO is positively skewed; in animals the median skewness is 3.37, with an interquartile range from 2.35 to 4.45. In plants, the median is 13.71 and the typical range is from 5.37 to 35.49. For comparison, the skewness of the exponential distribution is 2, which shows that individual stochasticity typically generates distributions of LRO that are more skewed than the exponential.

Kurtosis is seldom included in measures of variability, but it is important in these life cycles. Kurtosis is scaled relative to that of the normal distribution. The median kurtosis among animal populations is 15.27 (6.61 – 25.37), and among plant populations the median is 244.4 (38.86 – 1674.8). The tails of the distribution of LRO are thus far heavier than those of the normal distribution.

Plants life cycles create more individual stochasticity than do animal life cycles, and the ranges in variance, skewness, and kurtosis in LRO are greater among plant species than among animal species.

### 4.2 Longevity

In studies of longevity, coefficient of variation is preferred over standardized variance as a dimensionless measure of variation. The median CV of longevity in animals is 1.07 with typical values ranging from 0.88 to 1.25. In plants, the median is 1.26 with a range of typical values between 0.87 and 2.12.

Longevity is positively skewed, with rare individuals experiencing an unusually long life. In animals, the median skewness is 2.31 with an interquartile range from 1.99 to 3.08. In plants, the median skewness is 4.31 with typical values ranging from 2.34 to 9.16. These interquartile ranges suggest that distributions of longevity for plants are typically more skewed than for animals.

The median kurtosis in longevity among animals is 7.30 (4.32 – 11.10). In plants, the median kurtosis is 26.59 (8.01 – 133.86). The tails of the distribution of longevity thus appear to be heavier for plants than for animals.

### 4.3 Distributions across populations

Figure 1 shows the distribution of values of the various statistics of LRO, among populations of plants and animals (see Supplementary material for a similar graph for longevity). To make the patterns more clearly visible, for some statistics the data for plants and animals are trimmed by 20% (eliminating the top and bottom 10% of the values). See Tukey (1962) for a discussion of trimmed statistics.

**Figure 1:**
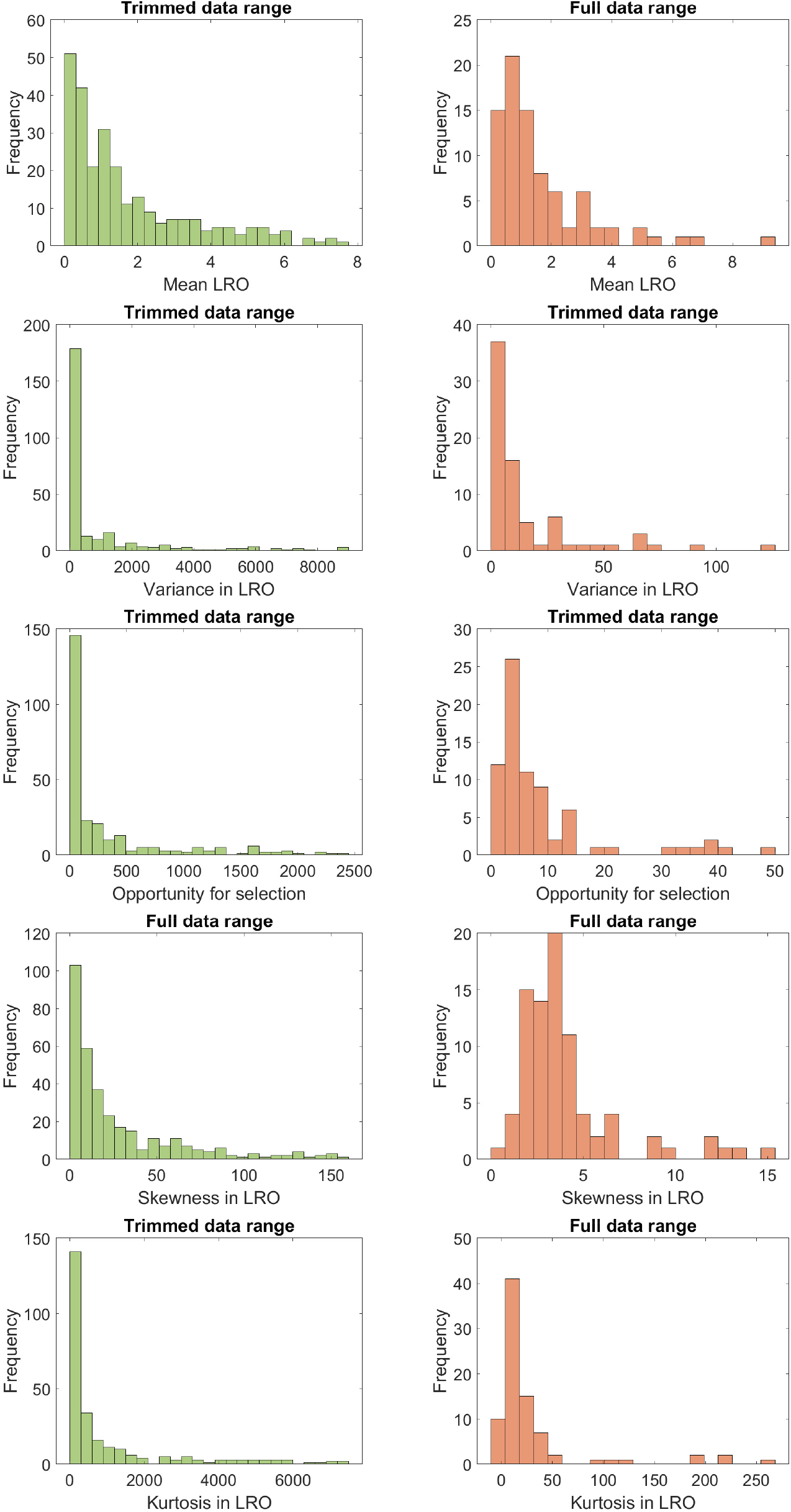
Histograms of the statistics of LRO (mean, variance, OFS, skewness, and kurtosis) for 332 populations of plants (left hand figures), and for 83 populations of animals (right hand figures). For both, some distributions were trimmed to better show the shape of the distribution; where the top of the plot reads “Trimmed data range” we left out 20% of the values for animal and plant populations (highest and lowest 10%).

Among the populations in our sample, plants exhibit a larger range of the statistics of LRO than animals and appear more skewed in the distributions of these statistics. In other words, plants show more uncertainty in their life history trajectories and the reproductive success they experience along the way than animals do. Additionally, plants, more so than animals, differ among populations in the level of this uncertainty.

## 5 Multidimensional life-history strategies

The various demographic outcomes do not vary independently among populations. We compute Pearson product-moment correlations. To do so, variables were log-transformed except for those that could take on zero or negative values, i.e. the kurtosis of LRO in animals, the kurtosis and skewness of longevity in both animals and plants, and the measures of variability of age at maturity for animals and plants. The correlations among the demographic outcomes are shown in Figure 5 and 6 in the Supplementary Material. Means and variances for the same life history trait are often strongly positively correlated. Skewness and kurtosis values are positively correlated with all other skewness and kurtosis values, and are negatively correlated to mean LRO and longevity. That is, longer-lived species, and species with high levels of reproduction tend to exhibit less reproductive uncertainty.

To move beyond pairwise correlations, we note that our analysis characterizes life histories by 16 demographic outcomes; these differ widely among plant and animal species. The life history of each species can be represented by a point in a 16-dimensional trait space. To reveal the patterns of association among traits, we used principal component analysis (PCA) to extract the major axes of variation. Principal component analysis reduces the dimension of the life history space into a set of orthogonal axes that best explain the variation among populations in these outcomes (Gaillard et al., 1989; Stearns, 1983; Salguero-Gómez et al., 2016).

Stearns (1983) applied principal component analyses to two sets of data on life history traits in 65 and 162 species of mammal, respectively, and concluded that their life histories are organized along an axis with fast life history strategies on one end and slow ones on the other (i.e. the fast-slow continuum). Gaillard et al. (1989) carried out a similar analysis on 80 mammal species and 114 bird species, which concluded that the variation of life history strategies of mammals and birds can be equally described with the fast-slow continuum. They found a secondary gradient corresponding to iteroparity. Salguero-Gómez et al. (2016) used PCA to analyze the life histories of 418 species of plants. They found two major axes of variation: an axis corresponding to the fast-slow continuum and one reflecting reproductive strategies. None of these analyses included measures of variance, skewness, or kurtosis in the set of life history traits.

We performed PCA analyses on the demographic outcomes after scaling them in such a way that the mean= 0 and SD= 1, as the demographic outcomes are expressed in different units.

### 5.1 Animals

The first three PC axes account for 36.4%, 28.2% and 13.2% of the variance, for a total of 77.8%. The loadings on each of the axes are given in Table 2. Figure 2 shows the distribution of animal species across the three axes, and the loadings of the most important demographic outcomes. The first axis separates species based on statistics related to uncertainty in LRO and longevity, with positive loadings for the opportunity for selection in LRO, and skewness and kurtosis in both LRO and longevity. High scores represent populations with high inter-individual variation relative to the mean of LRO, and a higher chance of extreme values in both LRO and longevity. We refer to this as the life cycle uncertainty axis.

**Table 2:** Loadings of lifetime reproductive output (LRO), longevity (LNG), generation time (GT), modes of parity (Parity), and age at maturity (AM) on the first three principal components of animals and plants.

| Demographic outcome | Animals |  |  | Plants |  |  |
| --- | --- | --- | --- | --- | --- | --- |
|  | PC1 | PC2 | PC3 | PC1 | PC2 | PC3 |
| LRO mean | -0.21 | 0.21 | -0.28 | 0.16 | 0.05 | 0.05 |
| LRO var | 0.08 | 0.15 | -0.36 | 0.10 | -0.15 | -0.14 |
| LRO OFS | **0.37** | 0.12 | -0.21 | -0.17 | **-0.44** | -0.02 |
| LRO skew | **0.39** | 0.14 | -0.08 | -0.17 | **-0.45** | -0.07 |
| LRO kurt | **0.37** | 0.18 | -0.09 | -0.17 | **-0.43** | -0.02 |
| LNG mean | -0.26 | 0.34 | -0.14 | 0.28 | 0.02 | 0.23 |
| LNG sd | -0.24 | **0.37** | -0.09 | **0.37** | -0.007 | 0.24 |
| LNG cv | 0.09 | 0.19 | **0.44** | 0.33 | -0.08 | -0.001 |
| LNG skew | **0.36** | 0.13 | -0.17 | -0.04 | -0.38 | 0.29 |
| LNG kurt | **0.35** | 0.15 | -0.23 | -0.08 | -0.35 | 0.34 |
| GT mean | -0.14 | **0.43** | 0.009 | **0.42** | -0.16 | 0.02 |
| GT sd | -0.16 | **0.41** | -0.04 | **0.41** | -0.13 | 0.16 |
| Parity | -0.16 | 0.11 | 0.11 | 0.12 | 0.03 | **0.51** |
| AM mean | 0.001 | 0.32 | 0.14 | 0.27 | -0.19 | **-0.43** |
| AM sd | 0.19 | 0.20 | **0.43** | 0.30 | -0.18 | **-0.39** |
| AM cv | 0.18 | 0.19 | **0.46** | 0.18 | -0.05 | -0.19 |

**Figure 2:**
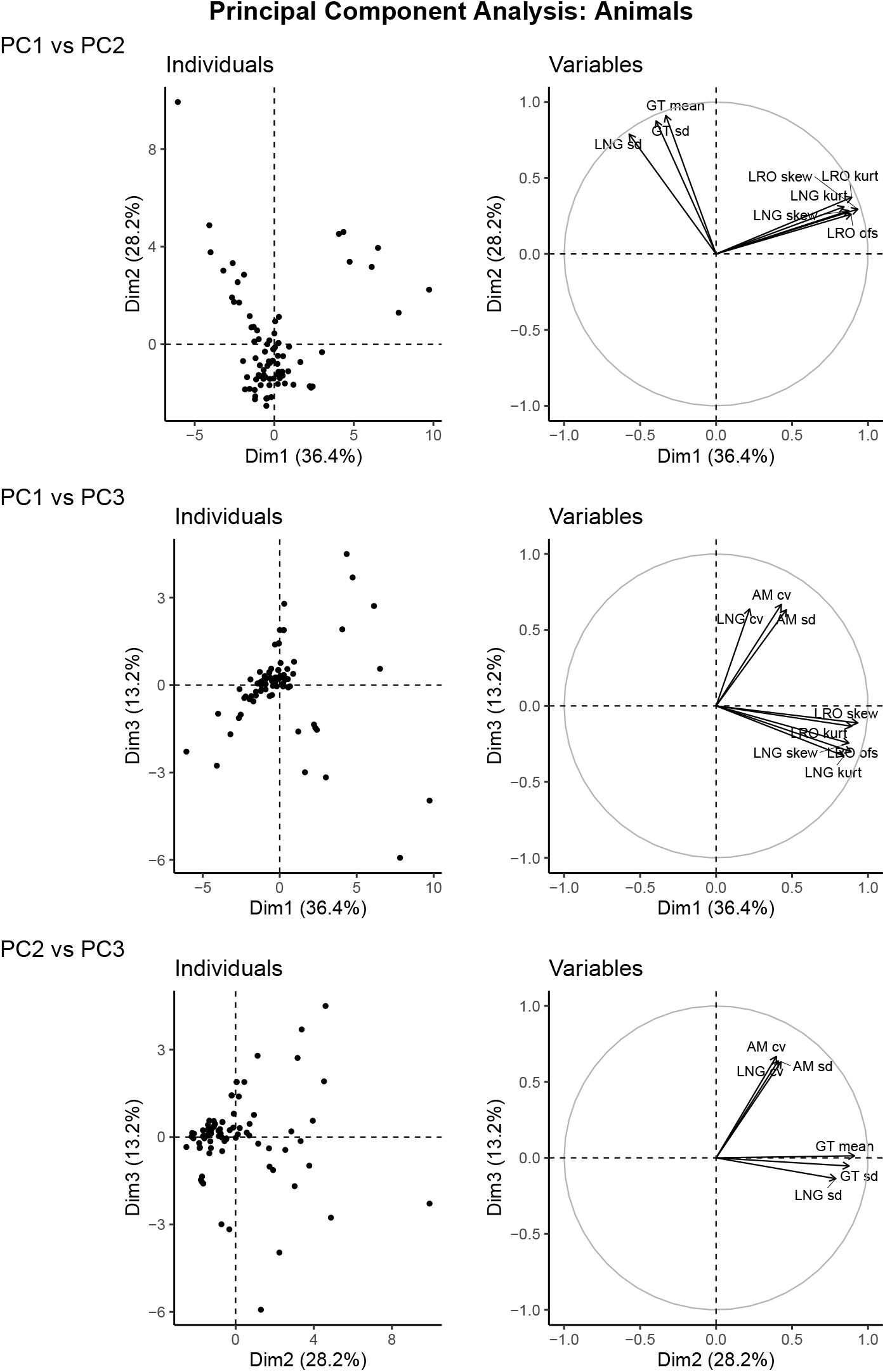
Principal component axes of 83 populations of animals. The right-hand graph shows the variables with the highest loadings onto these axes to facilitate visual understanding. Animal populations are primarily distributed along an axis representing measures of uncertainty in LRO and longevity (36.4%), followed by an axis representing generation time and longevity, and variation within these (28.2%). A third axis represents variation in timing of reproduction (13.2%).

The second axis captures variation in life cycle length. The mean and standard deviation of the age at offspring production (the mean of which represents generation time), as well as standard deviation in longevity, had the highest loadings on the second axis. We refer to this as the life cycle length axis.

The third axis separates populations on the basis of the timing of reproduction in the life cycle, with high loadings for the standard deviation of age at maturity, and the coefficients of variation in both age at maturity and longevity. Populations at the low end of this axis have low variation in the time to reproduction and time to death relative to the mean. We refer to this as the timing of reproduction axis.

### 5.2 Plants

The first three PCA axes for plants account for 28.2%, 22.9% and 11.9% of the variance, for a total of 63%. The loadings on each of the axes are given in Table 2. Figure 3 shows the distribution of 332 plant populations across three axes of variation, as well as the demographic outcomes with the highest loading on those axes. The first PCA axis captures life cycle length, with the highest loadings for generation time and standard deviation of the ages at production of offspring, and for the standard deviation of longevity.

**Figure 3:**
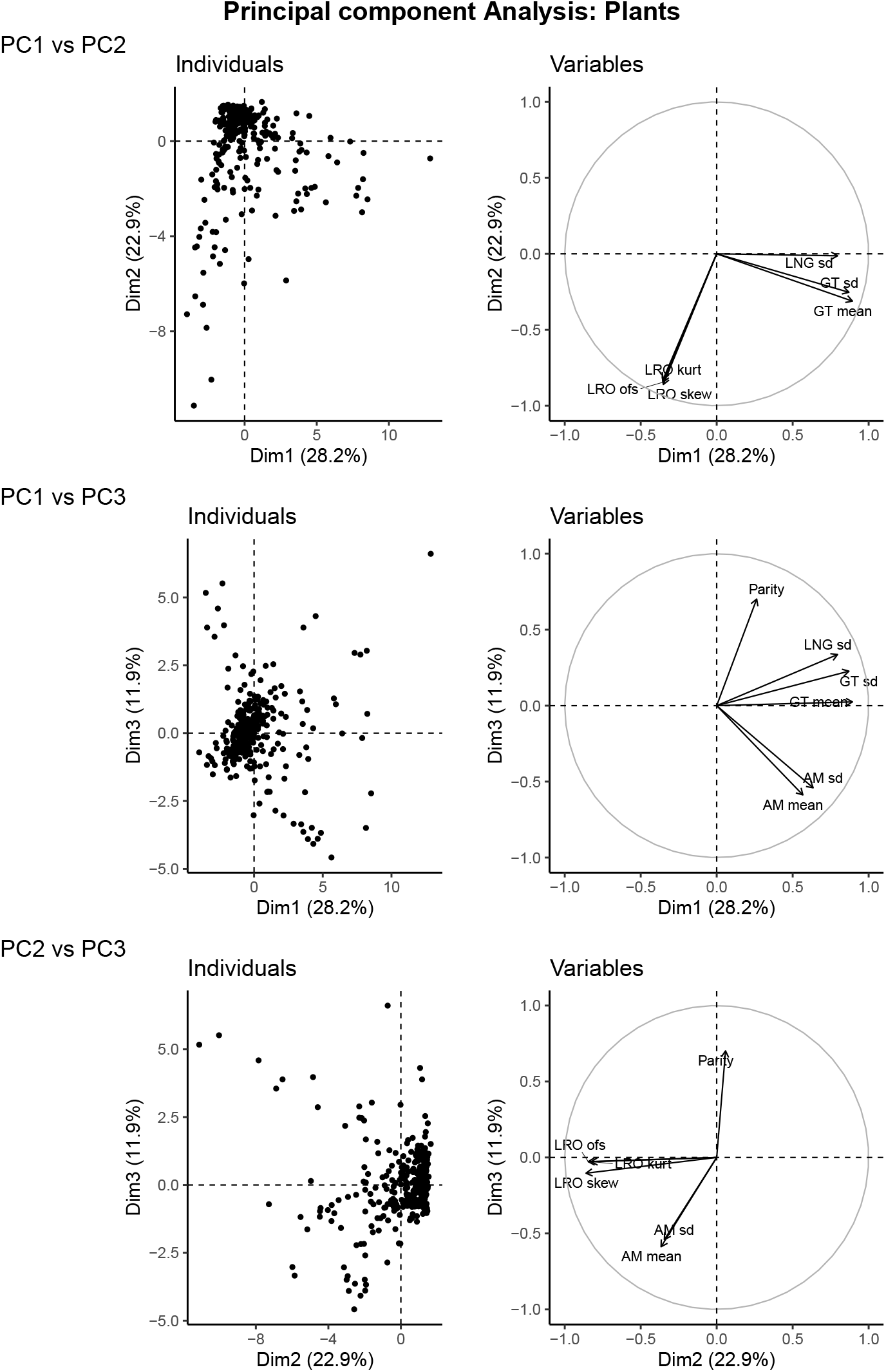
Principal component axes of 332 populations of plants. The right-hand graph shows the variables with the highest loadings onto these axes to facilitate visual understanding. Plant populations are primarily distributed along an axis representing generation time and longevity, and variation within these (28.2%), followed by an axis representing measures of uncertainty in LRO (22.9%). A third axis represents mean, variation, and spread of the timing of reproduction (11.9%).

The second PCA axis arranges species according to uncertainty in lifetime reproduction, with large negative loadings for the standardized variance, skewness, and kurtosis of LRO. At the negative end of this axis, populations have highly variable, skewed, and less predictable lifetime reproductive output.

The third PCA axis in plants is related to the spread of reproduction across the life cycle. Mean and standard deviation of age at maturity had high negative loadings, and mode of parity had a large positive loading on this axis. Populations at the positive end of this axis mature early and are iteroparous, whereas populations at the negative end of this axis have a shorter range of reproductive years, and mature later.

### 5.3 Comparisons of animals and plants

Animals and plants are distributed across similar life history axes. The first two axes reflect life cycle length and variance therein, and uncertainty in LRO. The differences lie in the order of the axes, and the fact that the uncertainty axis in animals also reflects uncertainty in longevity. The third axes both relate to timing of reproduction, but in animals this also correlates to variation in length of life, whereas in plants the mode of parity plays a larger role. In other words, in plants the third axis reflects the time available to reproduce relative to life cycle length, whereas in animals this axis captures how variable the timing of life cycle events is:

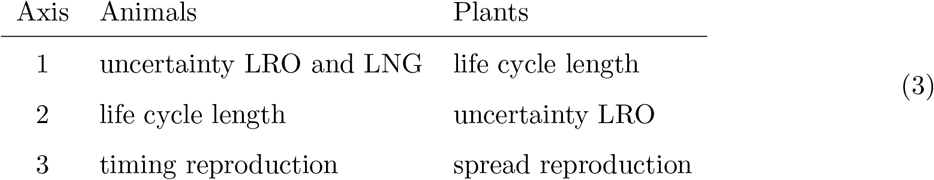

Measures of variation in demographic outcomes (variance, scaled variance, skewness, kurtosis) make important contributions to the axes along which life histories of both animals and plants are distributed. It is, arguably, no longer appropriate to ignore individual stochasticity in describing life histories.

## 6 Discussion

Individual stochasticity is a consequence of the life histories of all organisms. It generates variability among individuals in all demographic outcomes. This variability (measured, e.g., by variance, skewness, or kurtosis) is a consequence of the life cycle structure and demographic rates, just as population growth rate, stable stage distribution, the sensitivity and elasticity functions, and other familiar demographic outcomes are consequences. Given Markov chain and MCWR methods, it is now possible to calculate the variability among individuals implied by any life history.

To investigate the resulting patterns requires a sample of a variety of taxa. We have obtained such a sample from the Comadre and Compadre databases. These data are, in a sense, crowd-sourced information on life histories; as such, they provide an overview of the range of published demographic data for plants and animals. Although they are sourced from a scientific crowd (published studies), they are far from being a random or representative sample, either taxonomically or geographically (Salguero-Gómez et al., 2015, 2016). Even so, they give a picture of the range of what is possible.

The variability created by individual stochasticity is surprisingly large (Table 1, Figure 1). Plants are more variable than animals, in both longevity and LRO. This may reflect the greater flexibility in their life course due to their modular organization. Plant life cycles are often described in ways that provide more alternate pathways through the life cycle than is the case for animals. Shrinkage, for example, is common in plants, but less so in animals (Salguero-Gómez & Casper, 2010). Plants, perhaps more than frequently than animals, often experience extremely high mortality at early life history stages (seedlings). This naturally leads to high levels of variance in lifetimes among individuals.

The variability reported here is due to the individual stochasticity that is implied by the rates in the matrices **U** and **F**. Every individual experiences these rates and probabilities. In contrast, empirical measurements of lifetime demographic outcomes, obtained from longitudinal data on individuals followed through their lives, include both stochasticity and heterogeneity. Not every individual experiences the same rates, because they differ in factors not included in the demographic model. Thus the variance in the empirically measured outcomes reflects both stochasticity and whatever heterogeneity may exist among the individuals.

The standardized variance in LRO, which measures the opportunity for selection (Crow, 1958; Hersch & Phillips, 2004; Krakauer et al., 2011), has been measured empirically and provides a useful comparison to our studies. Robbins et al. (2011) reviewed empirical estimates of OFS for 22 species of birds and mammals. Their examples have a median of 0.55 and an interquartile range of 0.41–1.17. In 18 human populations, the range is even narrower, with a median of 0.34 and an interquartile range of 0.16–0.46 (Brown et al., 2009). This agrees rather well with the standardized variance calculated for a set of 40 developed countries analyzed by van Daalen & Caswell (2015, 2020b).

Our results clearly show more variance than these studies report, although our animal sample contains 3 populations within the IQR of Robbins et al. (2011). Whether this difference stems from the fact that these empirical studies mostly contain large mammals, or from issues with incorporating nulliparity in empirical studies (e.g., Klug et al., 2010; van Daalen & Caswell, 2020b), we cannot say. Potentially, the matrices we obtained lack certain processes that reduce inter-individual variation, although cases where additional population states reduce variance have not yet been reported^1^.

Stochasticity generates variance and uncertainty in life history outcomes such as LRO, longevity, age of reproductive maturity, and generation time. Beyond means, this study presented measures of variation and uncertainty in such life history traits for 332 plant and 83 animal populations. These measures of variability contribute to the life history strategy of a given population. Populations of plants and animals differ, not only in how fast they live, or what reproductive strategy they use, but in how flexible and uncertain their life histories are.

The surprising magnitude of the variation due to individual stochasticity, and its differences among species, cries out for its incorporation into the definition of the space within which life histories are distributed. This multidimensional space is defined by the principal component analyses axes. The first three axes account for 78% of the variance in animal, and 63% of the variance in plant, life histories. This is a step beyond the one-dimensional fast-slow life history continuum (Pianka, 1970; Stearns, 1983) which classifies life histories along a continuum from living fast, dying young and producing many offspring to living long and prospering by reproducing slowly but steadily. Later studies have added developmental rate (Stearns, 1983), iteroparity (Gaillard et al., 1989), and reproductive strategy (Salguero-Gómez et al., 2016; Paniw et al., 2018; Healy et al., 2019) as secondary axes of variation in life history strategy.

The life history strategies we identify reflect the inherent individual stochasticity in the life cycle. In both animals and plants, the first two axes resemble to some degree the fast-slow continuum and the reproductive strategies axis, but with the crucial distinction that these axes are dominated by inter-individual variation in demographic outcomes. The axis of life cycle length we find (first axis in plants, second axis in animals) incorporates the mean age at offspring production as generation time, and the standard deviations in age at offspring production and longevity. The uncertainty axis (second axis in plants, first in animals) reflects the standardized variance, skewness, and kurtosis in lifetime reproductive output. In animals, this axis furthermore relates to uncertainty in longevity.

Animal and plant populations differ, other than in percentage of variation explained in the first two axes, in a subtle way with regard to the third axis. The third axis, in both animals and plants, relates to the timing of reproduction in the life cycle. For animals, the third axis reflects variability in age at maturity, and standardized variability in longevity. In plants, the third axis reflects the start of the reproductive lifespan in mean and standard deviation of age at maturity, and the length of reproductive lifespan in mode of parity.

From these axes we ascertain that life history trade-offs and constraints are expressed not only in terms of means, but also in degrees of variation among individuals. Inter-individual variation is an important part of the life history strategy, and, therefore, cannot be ignored. This inter-individual variation, however, is completely due to individual stochasticity. Individual heterogeneity is another source of variation in life history outcomes.

If a putative source of heterogeneity is incorporated into the life cycle along with the original individual states, the variance calculated from the resulting model will reflect both sources. The variance in any outcome can then be decomposed into contributions of these sources of variance (Caswell et al., 2018). Studies that have done so generally find a large, or even overwhelming contribution of stochasticity. Only 5–10% of the variance in longevity in human populations could be attributed to heterogeneity in frailty (Hartemink et al., 2017) or socio-economic heterogeneity (Seaman et al., 2019). In other studies of longevity, a median of 35% of the variance could be attributed to unobserved heterogeneity in several laboratory studies of insects (Hartemink & Caswell, 2018). About 6% is due to unobserved heterogeneity in the Southern Fulmar (Jenouvrier et al., 2018).

Studies of variance components in lifetime reproductive output are rarer. Snyder & Ellner (2018) attributed about 39% of the variance in LRO to heterogeneous ‘quality’ in Kittiwakes. Jenouvrier et al. (2018) attributed 22% of the variance in LRO to unobserved heterogeneity in the Southern Fulmar. An analysis of the perennial herb *Lomatium bradshawii* in a stochastic fire environment found that the environment at birth could account for only 0.4% of the variance in LRO. (van Daalen & Caswell, 2020a). The broader life history effects of incorporating heterogeneity into models is yet unknown, as these multistate models are still rare.

Our selections from Comadre and Compadre include a wide range of species and life histories, from treecreepers to elephants, from herbs to trees. What if we treat these differences as an extreme case of heterogeneity? How much of the variance in longevity and LRO would be contributed by heterogeneity among populations of different species, in the face of the variance contributed by individual stochasticity within populations?

In general, the variance in some demographic outcome ***ξ*** is given by

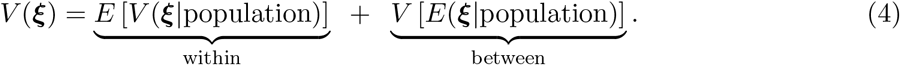

(e.g., Rényi, 1970; Frühwirth-Schnatter, 2006). The within-population variance is the contribution of individual stochasticity, since every individual within a population is subject to the vital rates of that population. The between-population variance is due to heterogeneity since it measures the extent to which the population differ in their rates. The contribution of heterogeneity is measured by the intraclass correlation coefficient (Falconer, 1960),

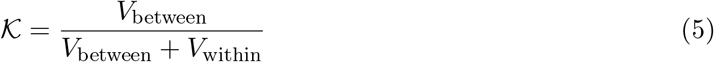

Applying this analysis to our results give, for longevity in animals

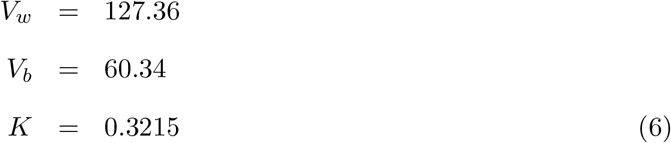

and for plants

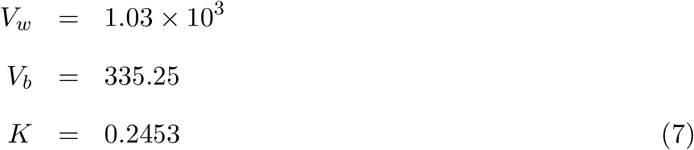

For LRO in animals

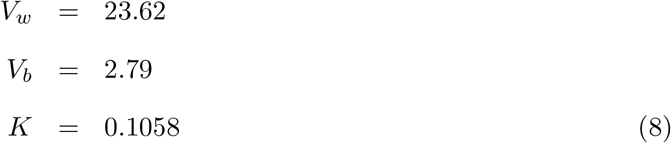

and for plants

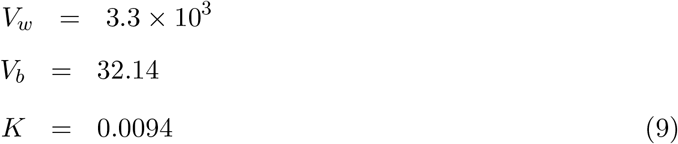

These results show that within-population variability due to individual stochasticity in movement through the life cycle overwhelms variation between populations. This is most apparent in the decomposition of variance in LRO in plants, where less than 1% of the variance is due to population identity, and 99% is due to individual stochasticity within populations. In animals, population identity underlies a greater part of the variance, but still only about 11%. Individual stochasticity determines 89% of the variation. The relative contribution of “heterogeneity” of course depends on both the mean and variance, with the variance between means giving the between-group component, and the mean of variances giving the within-group component. The difference between plants and animals then becomes clear; whereas plants and animals have a very similar median of mean LRO, plants have a 10-fold higher median of variance in LRO than animals (Table 1).

We conclude by mentioning a few extensions of the analysis here that could be applied in a comparative fashion. The analysis of LRO here can be applied to multiple kinds of offspring (Caswell, 2011). This would be particularly interesting as a comparison of sexual and clonal reproduction (we have not considered the latter) in plants. The comparisons among species could be informed by sensitivity analysis using results for LRO (van Daalen & Caswell, 2017), for longevity and generation time (Caswell, 2019), and the intraclass correlation coefficient 𝒦 (van Daalen & Caswell, 2020a). Here we have focused on the “mean” matrix reported in Compadre and Comadre. But it would be interesting to focus in more detail on the response of variability in demographic outcomes to environmental factors and stressors. Finally, the analysis here treats the matrices **U** and **F** as descriptions of a constant environment, but in cases where data are available, the calculations can be carried out in stochastic environments (Caswell, 2011; van Daalen & Caswell, 2020a).

## A Supplementary material

The calculations and analyses in this study were performed in R v3.5.3. The R package “*pracma*” was used to estimate the conditional number of matrices and, where necessary, pseudo-invert these in the calculations (Borchers, 2018).

### A.1 Measures of variance and uncertainty

The results of the calculations of life history outcomes are typically a set of vectors whose entries give the moments of some demographic outcome (***ξ***) for individuals starting in each stage. Let ***ξ***_*m*_ denote the vector for the *m*th moments. Then the variances are given by the vector

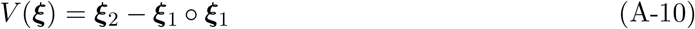

The skewness and kurtosis are most easily written in terms of the vectors of moments around the mean

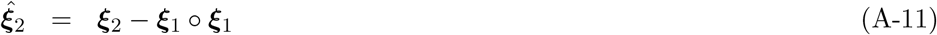

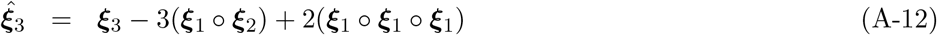

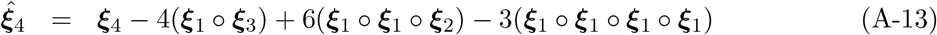

In terms of these central moments, the vectors giving the skewness and excess kurtosis of the elements of ***ξ*** are given by

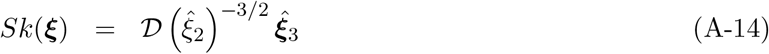

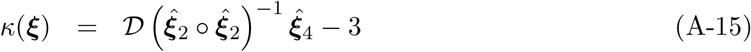

The skewness quantifies the asymmetry of the distribution; positive values indicate an extended positive tail, negative values indicate the opposite. Skewness can be interpreted as a measure of inequality. Kurtosis is a measure of the extent of extreme values, measured relative to the normal distribution. Positive kurtosis indicates a distribution with heavier tails (leptokurtic) and thus more likely to exhibit extreme values, either positive or negative.

### A.2 Calculation of individual stochasticity

The calculations of individual stochasticity rely on the Markov chain transition matrix **U** and the fundamental matrix

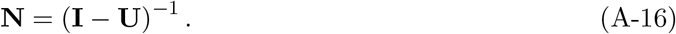

The (*i, j*) entry of **N** is the mean time spent in stage *i*, prior to absorption (i.e., death), of an individual starting in stage *j*.

#### A.2.1 Longevity

Longevity of an individual is calculated as the sum of the time spent in every transient stage, until eventual absorption (Caswell, 2009). The first four moments of longevity are given in Caswell (2013),

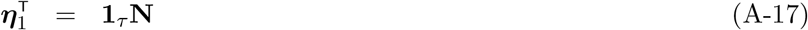

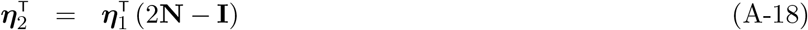

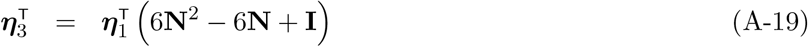

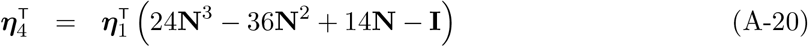

The *i*th entry of ***η***_*m*_ is the *m*th moment of longevity for an individual starting in stage *i*.

#### A.2.2 Lifetime reproductive output (LRO)

Calculation of LRO requires reward matrices whose entries give the moments of the reproductive output reward associated with each transition. Let **f** be a vector giving the mean reproductive output of each stage. If a single type of offspring is produced **f** ^T^ is the first row of the fertility matrix **F**. For species that produce multiple types of offspring (i.e., in which **F** contains positive entries in more than one row), we summed all types at each age and treated the sum as the mean stage-specific reproductive output. Modeling the number of offspring as a Poisson random variable with that mean gives the reward matrices

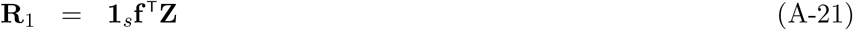

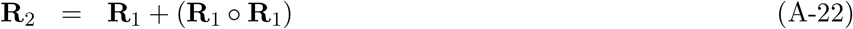

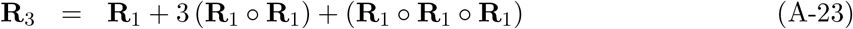

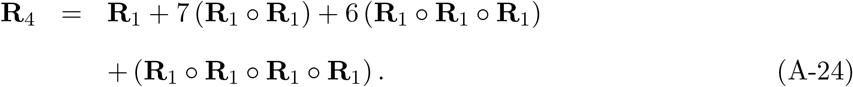

Following Theorem 1 of van Daalen & Caswell (2017), we write 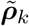 for the vector, of dimension *τ ×* 1 of the *k*th moments of lifetime reproduction for individuals starting in each transient stage. We define the matrix

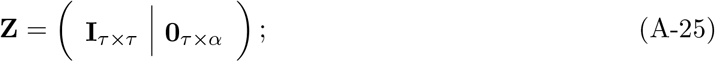

and also define 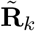, the *τ × τ* submatrix of **R**_*k*_ corresponding to transitions among the transient states:

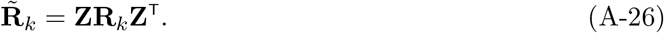

In terms of these quantities, the first four moments of LRO are

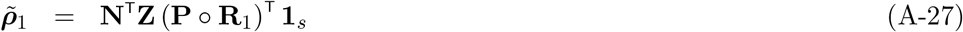

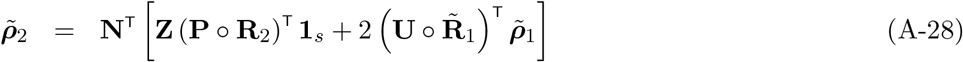

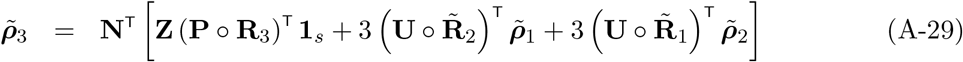

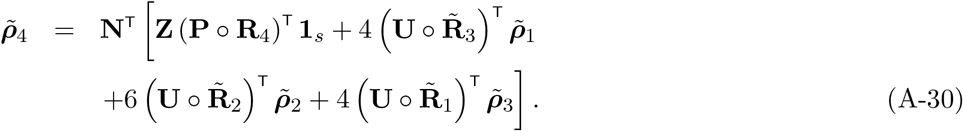

#### A.2.3 Age at maturity

The age at maturity is defined as the time to first enter any stage defined as reproductive, that is any stage *j* for which column *j* of **F** is non-zero. The technique, detailed in Caswell (2001, Section 5.3.3) has two steps. First, the transition matrix is modified to make the reproductive stages absorbing. An individual will end in one or the other of the two absorbing states, death-before-reproduction or reproduction-before-death. Then a conditional Markov chain is constructed, conditional on reaching reproduction. The age at maturity is the time to absorption in this conditional chain. We calculated the mean, standard deviation, and coefficient of variation of this time.

#### A.2.4 Generation time

The offspring production at age *x* of an individual starting in stage *j* is given by the vector

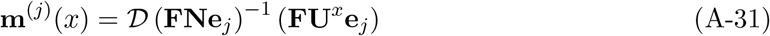

where the entries of **m**^(*j*)^ correspond to different types of offspring (Caswell, 2009).

The cohort generation time, given by the mean of this distribution, is

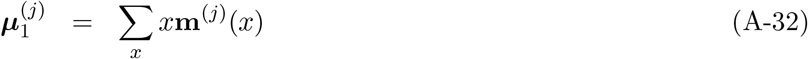

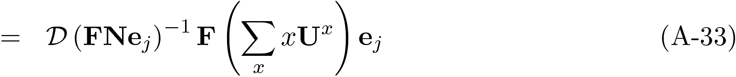

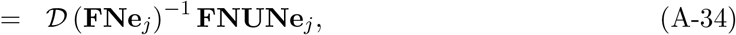

as in (Caswell, 2009).

#### A.2.5 Extent of iteroparity

We used the distribution **m**^(*j*)^ in (A-31) to derive a new result for the variance of the ages of mothers at the birth of offspring. The second moment of that age is

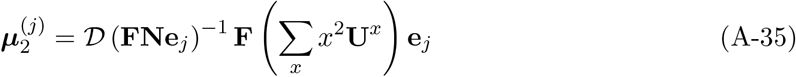

The new part of this expression is ∑_*x*_ *x*^2^**U**^*x*^. We can write and then simplify this summation,

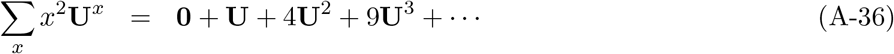

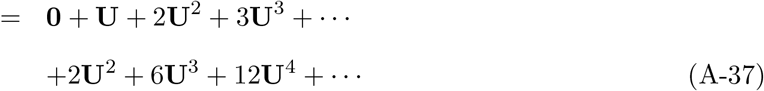

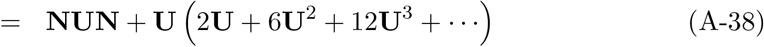

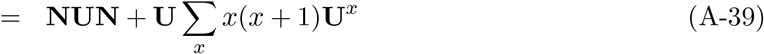

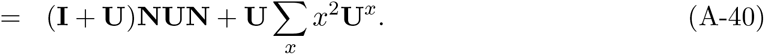

Solving for ∑_*x*_ *x*^2^**U**^*x*^, which appears on both sides, yields

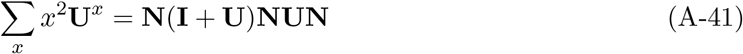

and thus

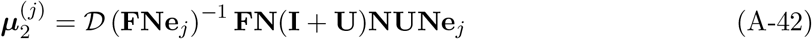

The vector of variances is given by

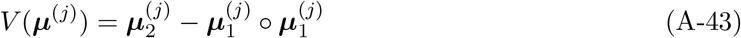

and from this we calculated the standard deviation and finally the coefficient of variation of the age of production of offspring. The coefficient of variation is dimensionless and is thus appropriate for comparing across life histories of different absolute lengths.

### A.3 Interspecific variation in statistics of longevity

Similar to the distributions for LRO in Figure 1, we show the distribution of the statistics of longevity for all plant and animal populations in Figure 4. We show trimmed histograms for some of the statistics for plants. Mean longevity, for example, has a high number of populations with a life expectancy of about 1. Paired with a long tail of extreme lifespans (up to a life expectancy of 150), the body of the distribution would be completely obscured if we showed the full range. The same is true for standard deviation, and kurtosis in longevity.

**Figure 4:**
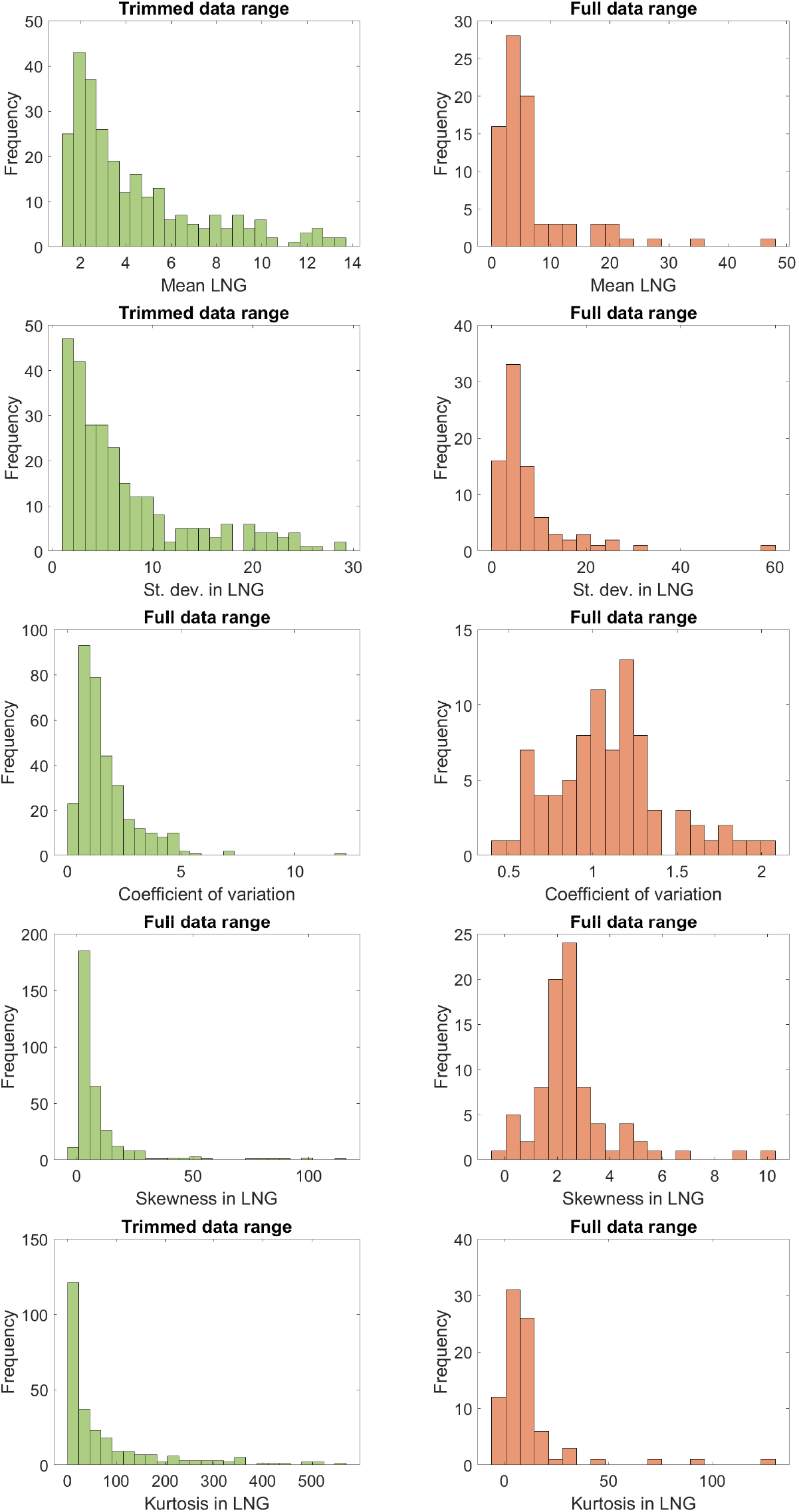
Histograms of the statistics of longevity (mean, standard deviation, coefficient of variation, skewness, and kurtosis) for 332 species of plants (left hand figures), and for 83 species of animals (right hand figures). For both, some distributions were trimmed to better show the shape of the distribution; where the top of the plot reads “Trimmed data range” we left out 20% of the values for animal and plant populations (highest and lowest 10%).

Plants show more variability among populations than animals do. They have a larger range in the values observed for each of the longevity statistics. All of these statistics indicate that plants experience greater variation and uncertainty in their life lifespans than animals do, although extreme examples can be found in both subsets.

### A.4 Correlations between life history outcomes

In figures 5 and 6, we show the Pearson product-moments correlations of all 16 of the demographic outcomes we include in our analyses for animals and plants, respectively. Some of these statistics are very tightly correlated; others not at all.

**Figure 5:**
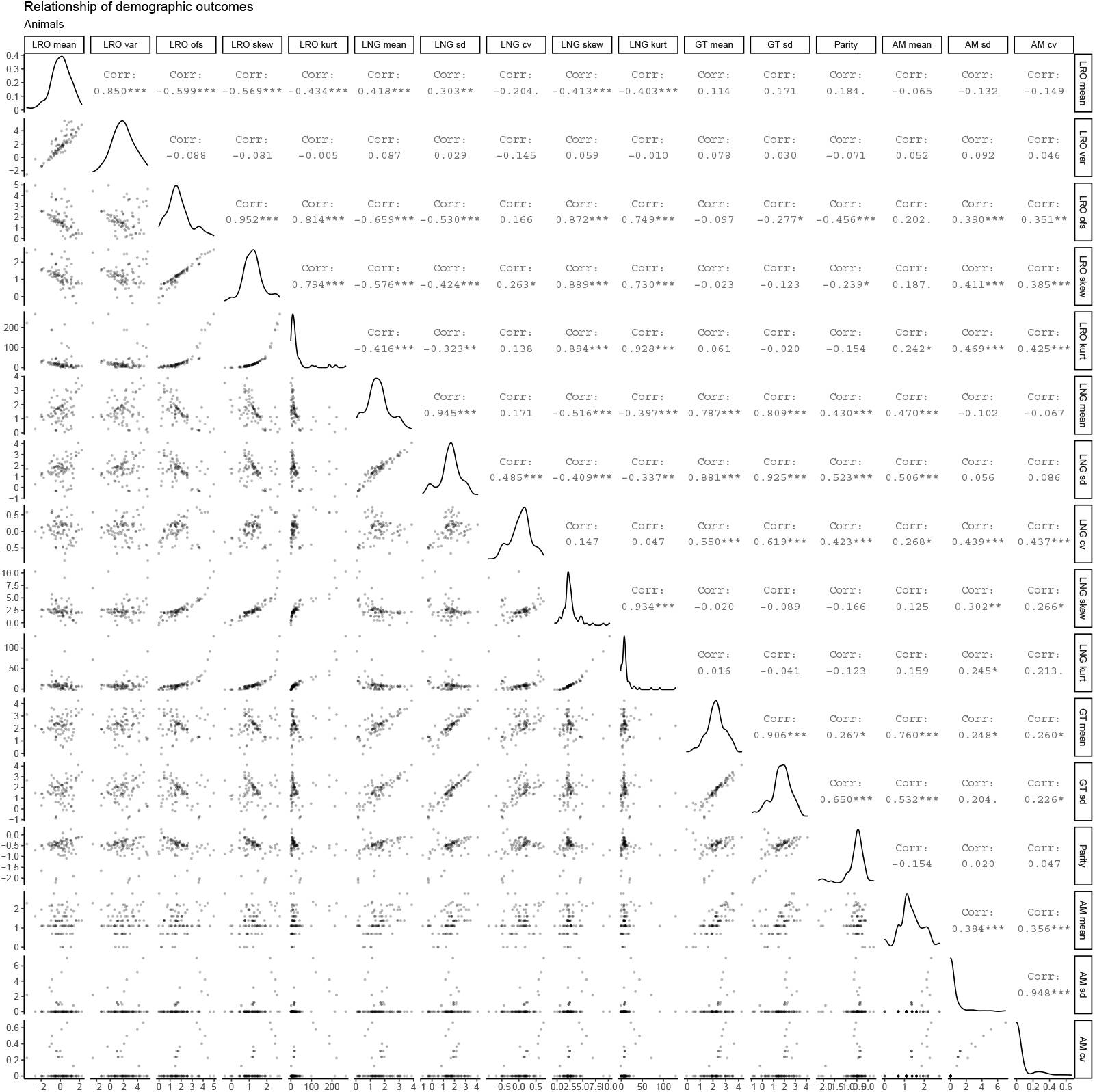
Relationship among the demographic outcomes in animals. Lifetime reproductive output (LRO), longevity (LNG), generation time (GT), modes of parity (Parity), and age at maturity (AM) were log-transformed, with the exception of the kurtosis of LRO, the kurtosis and skewness of longevity, and measures of variability in age at maturity. Pearson correlation coefficients are in the upper diagonal of the matrix while scatterplots are located in the lower diagonal. Finally, density plots are shown on the diagonal to visualize the distribution of the demographic outcomes.

**Figure 6:**
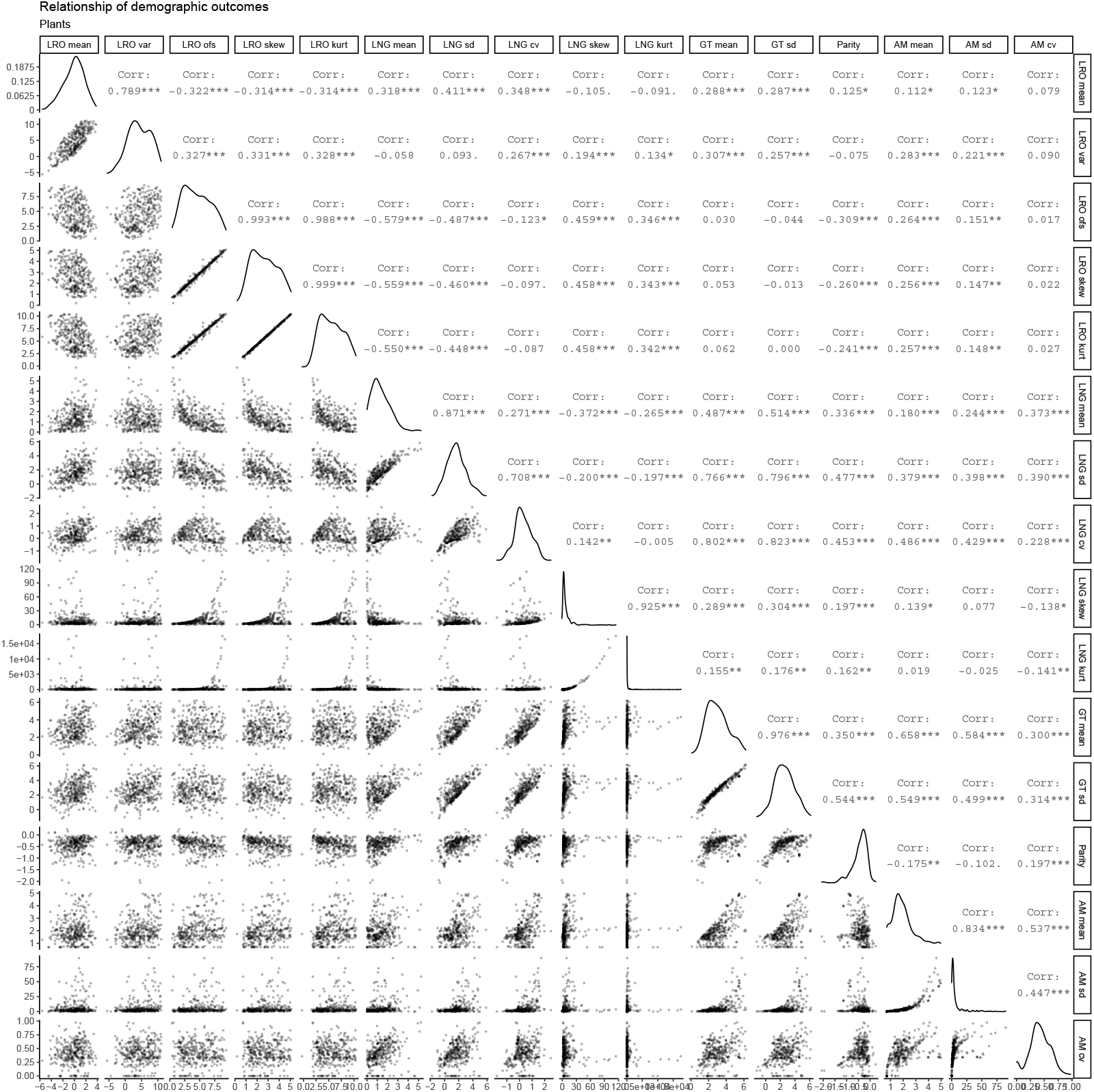
Relationship among the demographic outcomes in plants. Lifetime reproductive output (LRO), longevity (LNG), generation time (GT), modes of parity (Parity), and age at maturity (AFR) were log-transformed, with the exception of the kurtosis and skewness of longevity and measures of variability in age at maturity. Pearson correlation coefficients are in the upper diagonal of the matrix while scatterplots are located in the lower diagonal. Finally, density plots are shown on the diagonal to visualize the distribution of the demographic outcomes.

### Footnotes

1 There are theoretical examples that show additional structure in the population result in lower variance. A simple example would be variance in longevity in humans. If every individual has the same mortality rate *μ*, regardless of age (i.e. an alive-dead model), longevity has an exponential distribution of longevity with a standard deviation equal to the mean (1*/μ*). A typical human population with life expectancy of 70 years would show an SD of 70. But actual human populations, with strongly age-structured mortality, have a SD of longevity on the order of 10 years.

## Bibliography

Borchers, H. W. (2018). pracma: Practical Numerical Math Functions.

Brown, G. R., Laland, K. N., & Mulder, M. B. (2009). Bateman’s principles and human sex roles. Trends in Ecology & Evolution, 24 (6), 297–304.

Caswell, H. (2001). Matrix population models: Construction, analysis, and interpretation (2nd ed.). Sunderland: Sinauer Associates.

Caswell, H. (2009). Stage, age and individual stochasticity in demography. Oikos, 118, 1763–1782.

Caswell, H. (2011). Beyond R0 : Demographic Models for Variability of Lifetime Reproductive Output. PLoS ONE, 6 (6), 1–21.

Caswell, H. (2013). Sensitivity analysis of discrete Markov chains via matrix calculus. Linear Algebra and its Applications, 438 (4), 1727–1745.

Caswell, H. (2019). Sensitivity Analysis: Matrix Methods in Demography and Ecology. Springer Nature.

Caswell, H. (2020). The formal demography of kinship. II. Multistate models, parity, and sibship. bioRxiv.

Caswell, H., de Vries, C., Hartemink, N., Roth, G., & van Daalen, S. F. (2018). Age×stage-classified demographic analysis: a comprehensive approach. Ecological Monographs, 88 (4), 560–584.

Coale, A. J. (1972). The Growth and Structure of Human Populations: A Mathematical Approach. Princeton: Princeton University Press.

Cochran, M. E. & Ellner, S. (1992). Simple methods for calculating age-based life history parameters for stage-structured populations. Ecological Monographs, 62 (3), 345–364.

Crow, J. F. (1958). Some Possibilities for Measuring Selection Intensities in Man. Human Biology, 30 (1), 1–13.

Falconer, D. S. (1960). Introduction to quantitative genetics. New York: Ronald Press Co.

Frühwirth-Schnatter, S. (2006). Finite Mixture and Markov Switching Models. Springer Series in Statistics. New York: Springer-Verlag.

Gaillard, J.-M., Pontier, D., Allaine, D., Lebreton, J., Trouvilliez, J., & Clobert, J. (1989). An analysis of demographic tactics in birds and mammals. Oikos, 59–76.

Hartemink, N. & Caswell, H. (2018). Variance in animal longevity: contributions of heterogeneity and stochasticity. Population Ecology, 60 (1-2), 89–99.

Hartemink, N., Missov, T. I., & Caswell, H. (2017). Stochasticity, heterogeneity, and variance in longevity in human populations. Theoretical Population Biology, 114, 107–116.

Healy, K., Ezard, T. H., Jones, O. R., Salguero-Gómez, R., & Buckley, Y. M. (2019). Animal life history is shaped by the pace of life and the distribution of age-specific mortality and reproduction. Nature ecology & evolution, 3 (8), 1217–1224.

Hersch, E. I. & Phillips, P. C. (2004). Power and potential bias in field studies of natural selection. Evolution, 58 (3), 479–485.

Horvitz, C. C. & Tuljapurkar, S. (2008). Stage dynamics, period survival, and mortality plateaus. The American Naturalist, 172 (2), 203–215.

Jenouvrier, S., Aubry, L. M., Barbraud, C., Weimerskirch, H., & Caswell, H. (2018). Interacting effects of unobserved heterogeneity and individual stochasticity in the life history of the southern fulmar. Journal of Animal Ecology, 87 (1), 212–222.

Kimber, A. C. (1990). Exploratory data analysis for possibly censored data from skewed distributions. Journal of the Royal Statistical Society: Series C (Applied Statistics), 39 (1), 21–30.

Klug, H., Lindström, K., & Kokko, H. (2010). Who to include in measures of sexual selection is no trivial matter. Ecology Letters, 13 (9), 1094–1102.

Krakauer, A., Webster, M., Duval, E., Jones, A., & Shuster, S. M. (2011). The opportunity for sexual selection: not mismeasured, just misunderstood. Journal of evolutionary biology, 24 (9), 2064–2071.

Paniw, M., Ozgul, A., & Salguero-Gómez, R. (2018). Interactive life-history traits predict sensitivity of plants and animals to temporal autocorrelation. Ecology letters, 21 (2), 275–286.

Pianka, E. R. (1970). On r-and K-selection. The American Naturalist, 104 (940), 592–597.

Rényi, A. (1970). Probability Theory. Amsterdam: North-Holland.

Robbins, A. M., Stoinski, T., Fawcett, K., & Robbins, M. M. (2011). Lifetime reproductive success of female mountain gorillas. American journal of physical anthropology, 146 (4), 582–593.

Salguero-Gómez, R., Jones, O. R., Archer, C. R., Bein, C., de Buhr, H., Farack, C., Gottschalk, F., Hartmann, A., Henning, A., Hoppe, G., & others (2016). COMADRE: a global data base of animal demography. Journal of Animal Ecology, 85 (2), 371–384.

Salguero-Gómez, R., Jones, O. R., Archer, C. R., Buckley, Y. M., Che-Castaldo, J., Caswell, H., Hodgson, D., Scheuerlein, A., Conde, D. A., Brinks, E., & others (2015). The COMPADRE Plant Matrix Database: an open online repository for plant demography. Journal of Ecology, 103 (1), 202–218.

Salguero-Gómez, R., Jones, O. R., Jongejans, E., Blomberg, S. P., Hodgson, D. J., Mbeau-Ache, C., Zuidema, P. A., de Kroon, H., & Buckley, Y. M. (2016). Fast–slow continuum and reproductive strategies structure plant life-history variation worldwide. Proceedings of the National Academy of Sciences, 113 (1), 230–235.

Salguero-Gómez, R. & Casper, B. B. (2010). Keeping plant shrinkage in the demographic loop. Journal of Ecology, 98 (2), 312–323.

Seaman, R., Riffe, T., & Caswell, H. (2019). Changing contribution of area-level deprivation to total variance in age at death: a population-based decomposition analysis. BMJ Open, 9 (3), e024952.

Snyder, R. E. & Ellner, S. P. (2018). Pluck or Luck: Does Trait Variation or Chance Drive Variation in Lifetime Reproductive Success? The American Naturalist, 191 (4), E90–E107.

Stearns, S. C. (1983). The influence of size and phylogeny on patterns of covariation among life-history traits in the mammals. Oikos, 173–187.

Steiner, U. K. & Tuljapurkar, S. (2012). Neutral theory for life histories and individual variability in fitness components. PNAS, 109 (12), 4684–4689.

Steiner, U. K., Tuljapurkar, S., & Orzack, S. H. (2010). Dynamic heterogeneity and life history variability in the kittiwake. Journal of Animal Ecology, 79 (2), 436–444.

Steinsaltz, D. & Evans, S. N. (2004). Markov mortality models: implications of quasistationarity and varying initial distributions. Theoretical Population Biology, 65 (4), 319–337.

Tukey, J. W. (1962). The Future of Data Analysis. Annals of Mathematical Statistics, 33 (1), 1–67.

Tuljapurkar, S. & Horvitz, C. C. (2006). From stage to age in variable environments: life expectancy and survivorship. Ecology, 87 (6), 1497–1509.

Tuljapurkar, S., Steiner, U. K., & Orzack, S. H. (2009). Dynamic heterogeneity in life histories. Ecology Letters, 12 (1), 93–106.

van Daalen, S. & Caswell, H. (2015). Lifetime reproduction and the second demographic transition: Stochasticity and individual variation. Demographic Research, 33, 561–588.

van Daalen, S. & Caswell, H. (2020a). Variance as a life history outcome: Sensitivity analysis of the contributions of stochasticity and heterogeneity. Ecological Modelling, 417, 108856.

van Daalen, S. F. & Caswell, H. (2017). Lifetime reproductive output: individual stochasticity, variance, and sensitivity analysis. Theoretical Ecology, 10 (3), 355–374.

van Daalen, S. F. & Caswell, H. (2020b). Demographic sources of variation in fitness. In R. Sear, R. Lee, & O. F. Burger (Eds.), Human Evolutionary Demography. Open Book Publishers (in press).

